# Genomic signatures of local adaptation in recent invasive *Aedes aegypti* populations in California

**DOI:** 10.1101/2022.12.06.519335

**Authors:** Shaghayegh Soudi, Marc Crepeau, Travis C. Collier, Yoosook Lee, Anthony J. Cornel, Gregory C. Lanzaro

## Abstract

**Background:** Rapid adaptation to new environments can facilitate species invasions and range expansions. Understanding the mechanisms of adaptation used by invasive disease vectors in new regions has key implications for mitigating the prevalence and spread of vector-borne disease, although they remain relatively unexplored.

**Results:** Here, we use whole-genome sequencing data from 103 *Aedes aegypti* mosquitoes collected from various sites in southern and central California to infer the genetic structure of invasive populations. We integrate genome data with 25 topo-climate variables to investigate genome-wide signals of local adaptation among populations. Patterns of population structure, as inferred using principle components and admixture analysis, were consistent with three genetic clusters, likely resulting from multiple independent introductions. Using various landscape genomics approaches, which all remove the confounding effects of shared ancestry on correlations between genetic and environmental variation, we identified 112 genes showing strong signals of local environmental adaptation associated with one or more topo-climate factors. Some of them have known effects in climate adaptation, such as heat-shock proteins, which shows selective sweep and recent positive selection acting on these genomic regions.

**Conclusions:** Our results provide a genome wide perspective on the distribution of adaptive loci and lay the foundation for future work to understand how environmental adaptation in *Ae. aegypti* impacts the arboviral disease landscape and how such adaptation could help or hinder efforts at population control.

## Background

Biological invasions, involving the introduction, establishment, and spread of species outside their native zone, present one of the main threats to biodiversity, ecosystem integrity, agriculture, fisheries, and public health; with economic costs amounting to hundreds of billions of dollars per year worldwide (1, 2). During biological invasions, species often spread over a wide and climatically diverse range of environments. Although plasticity and broad ecological tolerance have been shown to facilitate the spread of invaders across such heterogeneous conditions (3, 4), increasing evidence suggests that rapid adaptation to local conditions is commonplace in invasive populations and can enable the establishment and spread of these species in the face of novel selection pressures (5–10). As such, invasive species represent an ideal model to investigate contemporary adaptive processes, which is key in an era of rapid, human-induced, environmental change.

The establishment and persistence of vectors within new ecological niches poses a serious threat from emerging and endemic arboviral diseases (11). Dengue fever is among the most widespread vector-borne infectious diseases in the world and is re-emerging in the United States of America after many years of absence (12, 13); the same trend is also reported elsewhere around the world (14). The risk of dengue infection coincides with the distribution of mosquitoes capable of transmitting dengue virus (DENV). *Aedes aegypti*, the yellow fever mosquito, is the primary urban vector of dengue viruses worldwide which is prevalent throughout the tropics and sub-tropics and is closely associated with human habitats outside its native range in Africa. The state of California has maintained an active and extensive mosquito surveillance program initiated in the early 1900s (15) and has previously only detected sporadic specimens of *Ae. aegypti* near airports (16). Confirmed breeding populations of *Ae. aegypti* in California were never reported prior to the summer of 2013, when they were detected in three cities in the central valley counties of Fresno and Madera and the coastal county of San Mateo (16, 17). Subsequent reports indicate that *Ae. aegypti* has now become established and is spreading throughout large regions of California (18). Recent studies demonstrated that populations of *Ae. aegypti* in California were presumably introduced from multiple, genetically distinct source populations in the U.S. and/or northern Mexico (18, 19). These populations have also undergone behavioural and genetic changes in comparison to their ancestral African form, including the evolution of house-entering behaviour and a preference for human odour and blood-feeding (20, 21).

Although it is known that the environment is a key element in driving and altering the life-history traits of *Aedes* mosquitoes (22–24), there remains a limited understanding of how their genomic background changes across a heterogeneous landscape. A landscape genomics approach is an important first step to associate population structure with the environment and to narrow down candidate genomic targets for further investigation of local environmental adaptation (25). In the present study, we first investigated the genetic structure of *Ae. aegypti* populations distributed in southern and central California and then applied landscape genomics approaches to test the possibility of rapid adaptation to heterogeneous environments by identifying loci with unusual allelic associations to different environmental conditions. We produced evidence relevant to the question of whether adaptation is predominantly mono- or polygenic by conducting genotype-environment association (EAA) analysis using whole genome re-sequencing (WGS) data and by characterizing population structure to account for potentially confounding effects in EAA tests. Our new insights into the evolution of rapid adaptation observed in *Ae. aegypti* in California will improve our knowledge of evolutionary forces and processes during the invasion of disease vectors, which is crucial for advancing dynamic mitigation strategies aimed at reducing disease risk worldwide (26, 27).

## Materials and Methods

### Mosquito collections

A total of 103 individual adult female *Ae. aegypti* from 12 populations were collected from 33 locations between 2013-2017 (Figure 1, Supplementary Table 1). These mosquitoes were collected using BG Sentinel traps baited with CO_2_. All collections on private properties were conducted after obtaining permission from residents and/or owners. Mosquito samples were individually preserved in 80% ethanol and held at either −20 or −80°C prior to DNA extraction.

**Figure 1.**
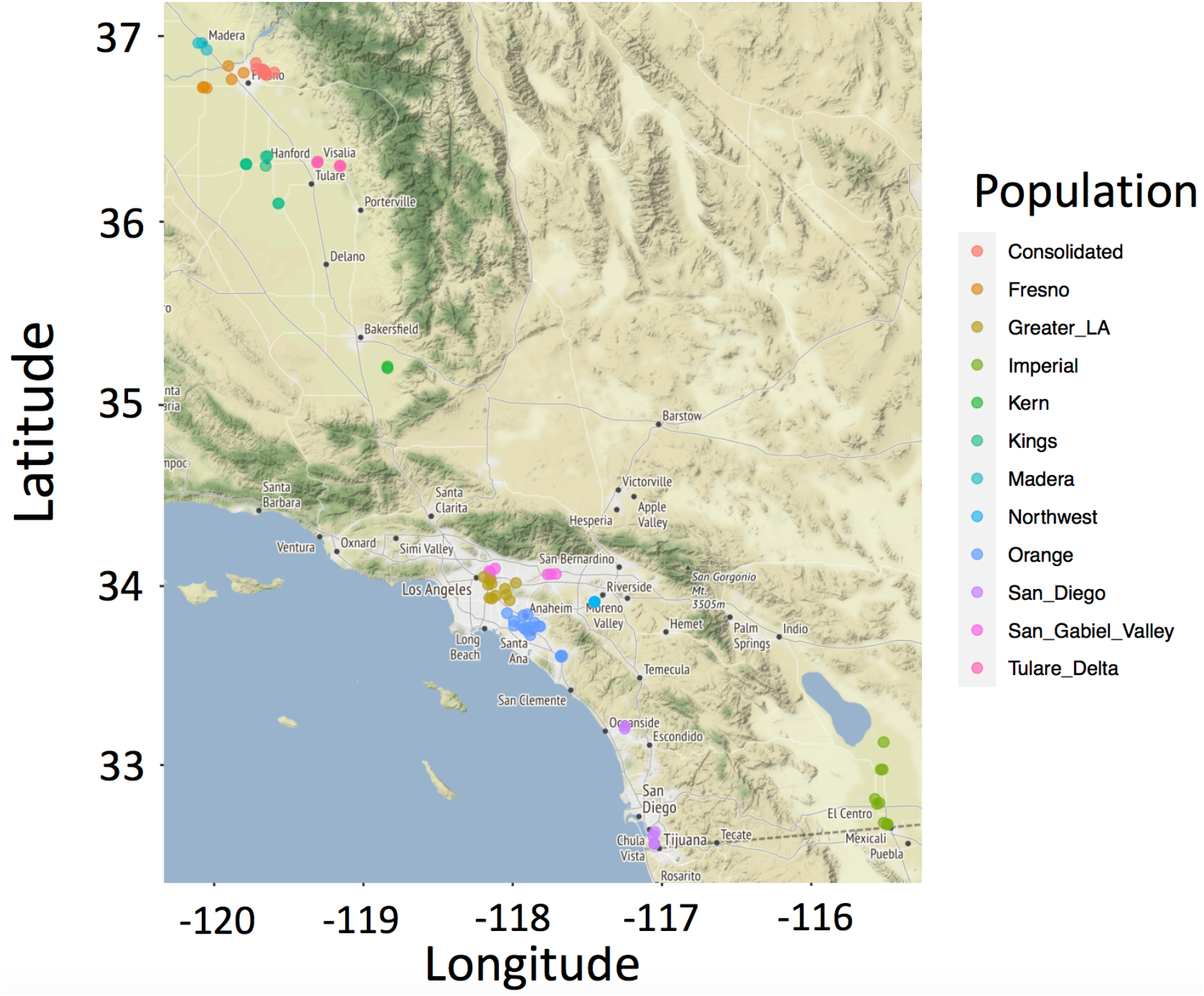
Sampling locations of 103 Ae. aegypti mosquitoes collected across central and southern California between 2013 and 2017. Map was created using R Project for Statistical Computing v. 3.3.1 (28) and package maps v. 3.2.0 (29). Colors indicate the origin mosquito abatement or vector control district of populations.

### Whole-genome resequencing

Genomic DNA was extracted and sequenced using established protocols as described by Nieman et al. 2015 (30). Genomic DNA concentrations for each sample were quantified using the Qubit dsDNA HS Assay Kit (Life Technologies, Carlsbad, CA) on a Qubit instrument (Life Technologies, Carlsbad, CA). A genomic DNA library was constructed for each individual mosquito using 20 ng DNA, the Qiaseq FX 96 kit (Qiagen, Valencia, CA), and Ampure SPRI beads (Beckman Coulter, Brea, CA) following an established protocol by Nieman et al. 2015 (30). Library concentrations were measured using Qubit (Life Technologies, Carlsbad, CA) as described above. Libraries were sequenced as 150-bp pair-end reads, each in one lane of an Illumina HiSeq 4000 platform at the UC Davis DNA Technologies Core and according to the manufacturer’s standard protocols (summary statistics of Illumina re-sequencing data per sample is available in Supplementary Table 1).

### Alignment, variation calling and annotation

Raw reads were trimmed using Trimmomatic (31) version 0.36 and high-quality trimmed reads were mapped to the AaegL5 reference genome (32) using BWA-MEM version 0.7.15 with default parameters. Mapping statistics were calculated using Qualimap (33) version 2.2 (Supplementary Table 1). The marked duplicate reads were removed using Picard tools version 2.1.1 (http://broadinstitute.github.io/picard/).

We called variants using Freebayes (34) version 1.0.1 with standard filters and population priors disabled. We required a minimum depth of 8 to call variants for each individual following the recommendation of Crawford and Lazzaro to minimize bias in population inference (35). To improve the reliability of calls, we required variants to be supported by both forward and reverse reads overlapping the loci (Erik Garrison, Welcome Trust Sanger Institute and Cambridge University, personal communication, Dec. 2014). The repeat regions were “soft-masked” in the AaegL5 reference genome and single nucleotide polymorphisms (SNPs) in these regions were excluded from analysis. SNPs with minor allele frequency (MAF) of < 3% and individuals with > 20% missing genotypes were excluded from the analysis to minimize bias from sequencing error (22).

### Analysis of population structure

We started by generating linkage disequilibrium (LD) pruned SNP sets as follows. We set sliding widows of size 50 (that is the number of markers used for linkage disequilibrium testing at a time) and window increments of 5 markers. For any pair of SNPs in a window we defined, the first marker of the pair was discarded when the correlation coefficient (r^2^) between markers exceeded 0.2 using an R package, SNPRelate (36). This yielded 100,089 independent SNPs that were retained for downstream population structure analysis.

Analysis of population structure was performed using the quality-control-positive linkage-disequilibrium-pruned set of 100,089 autosomal SNPs. Principle component analysis (PCA) (37) was conducted across all populations using EIGENSTART (v. 6.1.4) and results were visualized in RStudio (38). We applied unsupervised hierarchal clustering of individuals using the maximum likelihood method implemented in ADMIXTURE (v. 1.3.0) (39) using default input parameters. ADMIXTURE estimates ancestry coefficients from K modelled ancestral populations by assigning individuals to subpopulations after maximizing Hardy-Weinberg equilibrium of allele frequencies. The ‘—cv’ flag was added to perform the cross-validation procedure and to calculate the optimal number of K. A good value of K exhibits a low cross-validation error compared to other K values.

### Environmental data

A total of 25 biologically relevant topo-climate variables (Supplementary Table 2) were used in the analyses. Climate data for each population were collected from geographic coordinates of the locations where the samples were collected using the software package ClimateNA (40). All available data points from 2010 to 2017 were used.

### Screening for SNPs associated with local adaptation

To identify putative loci with a signal of selection, we used three approaches with different underlying algorithms and assumptions. To identify loci associated with environmental predictors two EAA approaches, BayPass (41) and latent factor mixed model (LFMM) (42), were implemented.

BayPass package version 2.1 (43) provides a re-implementation of the Baysian hierarchical model and explicitly accounts for the covariance structure among population allele frequencies that arises from the shared populations history. This was achieved by estimating a population covariance matrix, which renders the identification of SNPs subjected to selection less sensitive to the confounding impact of neutral genetic structure (41). Population structure was estimated by choosing a random and unlinked set of 10K SNPs using the BayPass core model. We used the AUX covariate model to assess the association of SNPs with topo-climate l variables. For each SNP, the Bayes factor (denoted BF_is_ as in Gautier, 2015 (43),) relies on the importance sampling algorithm proposed by Coop et al. 2010 (44) and uses Markov Chain Monte Carlo (MCMC) samples obtained under the core model. BF_is_ was expressed in deciban units (db) via the transformation 10 log_10_ (BF). As a decision rule and to calculate a significance threshold for outlier identification, pseudo-observed data (POD) were employed with the same random 10kb SNPs used for the core model, and a 1% empirical threshold was calculated for the observed Bayes factor. To produce a narrower set of outlier loci, we then followed the popular Jeffreys’ rule (45) that identified outlier loci with BF ≥ 10. The Latent Factor Mixed Model (LFMM) is a variant of the Bayesian principal component analysis in which residual background population structure is introduced via latent factors. We used a model with three latent factors to account for neutral population structure in the data based on the result we obtained from PCA and ADMIXTURE. We ran 10^5^ MCMC integrations with 5 burn-in steps with 10 replicate runs. Z-scores from replicate runs were combined and adjusted using the genomic inflation factor which estimates the excess of the false discovery rate due to multiple testing, and it is defined as the ratio of the observed and the expected median of the distribution of the test statistic (46). Lambda was calulated as median (Z^2^)/0.456. We corrected for multiple testing by fixing the false discovery rate to 5%. Only SNPs with FDR < 5% were retained as those significantly associated with topo-climate variable.

In addition to two EAAs methods, PCAdapt was used to find loci putatively under selection pressure as they deviate from the typical distribution of the test statistic Z (47). Similar to LFMM, three K populations were chosen to account for neutral population structure. PCAdapt examines the correlations (measured as the squared loadings *p^2^_jk_*, which is the squared correlation between the *j*th SNP and *K*th principal component) between genetic variants and specific PCs without any prior definition of populations. Assuming a chi-square distribution (degree of freedom = 1) for the squared loadings *p^2^_j1_*, as suggested by Luu et al. 2017 (47), we used PCAdapt to calculate *P* values for all SNPs and then estimated the FDR to generate a list of candidate SNPs showing significant associations to population structure. Only SNPs with FDR < 5% were retained as those significantly involved in local adaptation.

### Identification of top candidate genes

Loci that selected as outliers by all three implemented methods, BayPass, LFMM and PCAdapt, were identified. For each gene that we annotated in the genome we counted the number of outlier SNPs f (a) and the total number of SNPs (n). To identify top-candidate genes for each variable, we compared the number of outlier SNPs per gene to the 0.9999 quantile of the binomial expectation where the expected frequency of SNPs per gene is *p* = ∑ *ai/ni* (summation over *i* genes), calculating *p* separately for each environmental variable and excluding genes with no outliers from the calculation of *p*. Any genes with *p* values falling above this cutoff threshold were then identified as “top candidate genes” (48). The position and function of the candidate genes identified by this approach were mined using the mosquito genomics resource of VectorBase (49).

### Signature of positive selection around candidate genes

Two standard methods were further applied to search for signs of selective sweep in different groups of populations. Pairwise nucleotide diversity (π) (50), which is expected to have local reduction following a selective sweep, was calculated using a sliding window approach with window size of 10kbp and moving step of 5kbp using the software R package PopGenome (51) separately for each of the three groups detected by the PCA and admixture analyses. Weir and Cockerham’s Fst, which measures genetic divergence between pairs of three groups of populations, was calculated using a sliding-window size of 10kb and moving step of 5kb by VCFtools (52).

### Gene annotation and enrichment analysis

To explore which biological processes (BP) top candidate genes are involved in, we performed a Gene Ontology (GO) and enrichment analysis for the top candidate genes we identified using topGO package in R (53). Significance for each individual GO-identifier was computed with Fisher’s exact test and significance threshold of 1%. We also performed BLAST (54) searches of the predicted genes against the homologous genes in the annotated *Drosophila melanogaster* genome in order to potentially obtain more precise information on their functional annotation.

## Results

### Characterization of sequence variation in *Ae. aegypti*

*We* performed whole genome re-sequencing of all 103 *Ae. aegypti* samples and obtained, on average, over 110 million Illumina raw reads with an average sequencing depth of ~10X per individual covering > 85% of the reference genome. After variant calling and applying appropriate filtering, we identified a total of 1,968,198 single nucleotide polymorphisms (SNPs) with a minor allele frequency (MAF) > 3% which were subjected to downstream analysis. Supplementary Table 1 summarizes the per-individual read counts and coverage depths.

### Analysis of local population structure

We examined population structure and identified ancestral components with an autosomal marker dataset (100,089) using PCA and ADMIXTURE. We found a strong local population structure across the entire range of collections by PCA. The two-first axes (principal components 1 and 2) explained a large proportion of the variation, cumulatively accounting for 63.9% of the variance in SNP genotypes and three main genetic clusters were determined from this analysis (Figure 2a). The first cluster (*A.ae1*) included samples collected from various sites in Consolidated Mosquito Abatement District in central California. The second cluster (*A.ae2*) primarily included populations from other mosquito districts in central California (Madera, Fresno, Kings, Tulare_Delta, and Kern) and the third cluster (*A.ae3*) consisted of populations from southern California mosquito districts (San_Diego, Imperial, Orange, Northwest, Greater_LA, and San_Gabriel_Valley).

**Figure 2.**
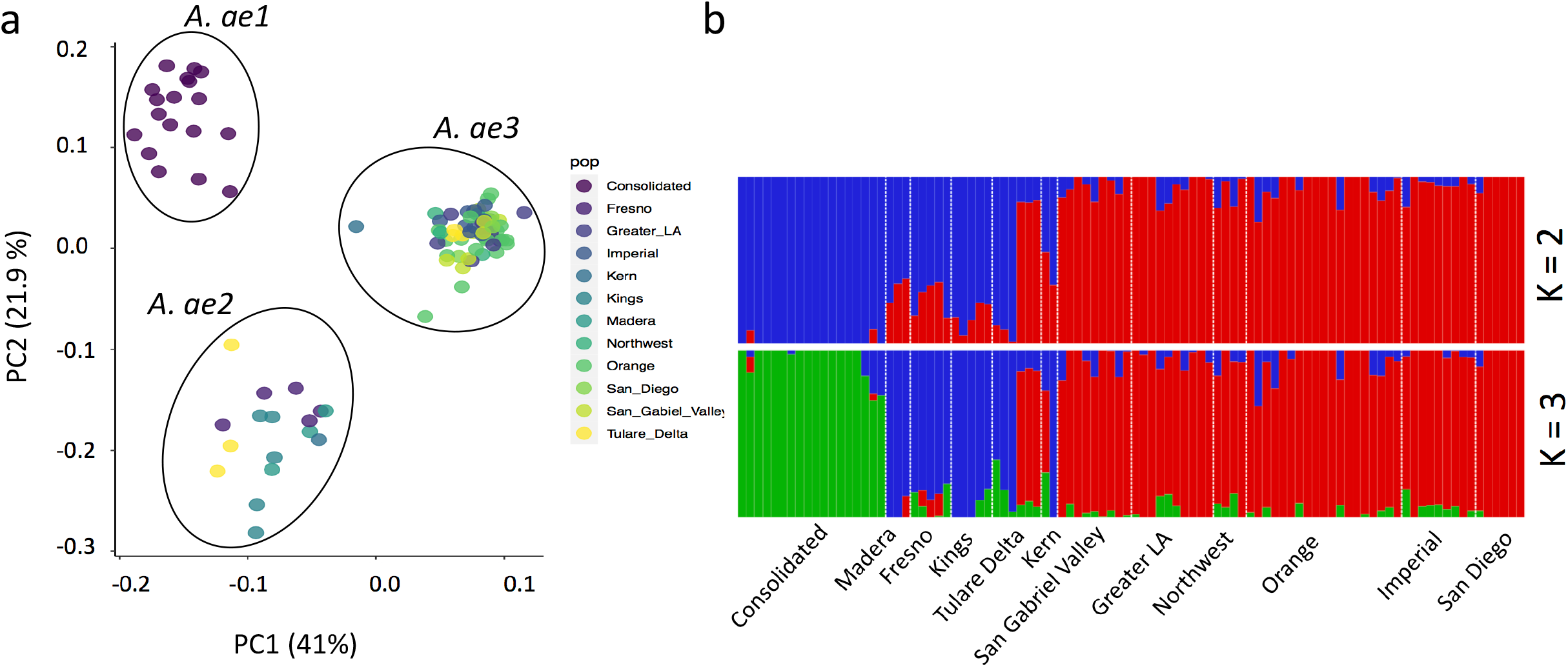
**a.** The first two principal components of a principal component analysis (PCA) of individual genotypes based on the LD pruned dataset describing the relationship among populations. Color code refers to origin mosquito abatement or vector control district of populations. **b**. Clustering assignments of each genotype inferred using the software Admixture for K = 2 and K = 3 populations. Each color represents one genetics cluster and each vertical bar represents one genotype.

Admixture analysis highlighted a significant population structure. According to cross validation error (Supplementary Figure 1), k=3 was the most well-presented population structure for our dataset which distinguished individuals from southern California, central California, and Consolidated as genetically distinct groups (Figure 2b). There were some individuals positioned between the three main clusters suggesting a potential admixture between different populations (Figure 2a/b). Our results generally recapitulate the broad inferences of a previous study by Lee et al 2019 (18).

### Genomic evidence for local adaptation in response to environmental heterogeneity in California

If these mosquitoes were locally adapted, we would expect to see that these populations of *Ae. aegypti* harbor genomic loci with signals of selection correlated to heterogeneous environmental conditions after taking the underlying population structure into account. In order to find genomic regions that are associated with local adaptation and to assess how candidate variation is portioned among different environmental variables, we carried out three complementary approaches which take into account the neutral genetic structure.

We performed PCA analysis for the 25 topo-climate variables extracted from ClimateNA (Supplementary Table 2). The two-first axes (PC1 and PC2) explained a large proportion of the variation, 56% and 35% respectively. Twelve *Ae. aegypti* populations, mainly distributed along the second PC axis, were linked to both temperature and precipitation variables (Supplementary Figure 2). We then started by identifying SNPs that showed strong associations with the topo-climate variables using LFMM and BayPass (42, 43). The number of latent factors was set to three based on the results of Admixture and PCA, as explained above. Under K = 3 genetic clusters, LFMM identified 17,519 outlier SNPs with a genomic signal of local adaptation at the FDR of 5% across all variables. Among all variables, we found the highest number of outliers associated with both temperature and humidity (climatic moisture deficit, degree-days above 18°C, and annual heat-moisture index with 4,406; 4,078; and 3,685 outlier SNPs respectively).

LFMM is robust in identifying adaptive processes that result from weak, multi-locus effects across various demographic scenarios and sampling schemes. However, it is important to recognize that a subset of the 17,519 candidate loci identified through this single analysis are likely to be false positives. We therefore explored associations with the Bayesian method available in BayPass under the AUX covariate model. We selected this model over others because it is more precise and efficient when estimating the covariance matrix (Ω) and more sensitive for identifying SNPs displaying weak association signals resulting from soft selective sweeps often involved in polygenic characters (43). Analysis of the data set under the BayPass core model allowed us to estimate the scaled covariance matrix of population allele frequencies Ω that quantifies the genetic relationship among each pair of populations. The resulting estimates of Ω accurately reflected the known structure between samples, that is, a clustering at the higher level by population geographic origin (Supplementary Figure 3a and 3b). BayPass analysis identified 16,976 SNPs with a signature of selection widespread across the genome and associated with various topo-climatic factors we tested. Among the analyzed variables, latitude, annual heat-moisture index, mean annual temperature, and climatic moisture deficit were the variables with the highest number of outlier SNPs detected by the BayPass AUX model.

The PCAdapt (47) method is considered less sensitive to confounding demography due to its ability to account for population structure or unobserved spatial autocorrelation in the data (55). Compared to 17,519 and 16,976 outlier SNPs detected by LFMM and BayPass respectively, PCAdapt identified a of total of 8,637 SNPs with a signature of selection widespread across the genome. Figure 3 shows an example of a circular Manhattan plot for a single environmental variable: mean warmest month temperature (MWMT). Across all three implemented methods, there were 1,991 SNPs consistently identified as outliers with a signal of selection and correlated with topo-climate variables, providing higher confidence that these loci are located within, or close to, regions involved in adaptation to heterogeneous environments.

**Figure 3.**
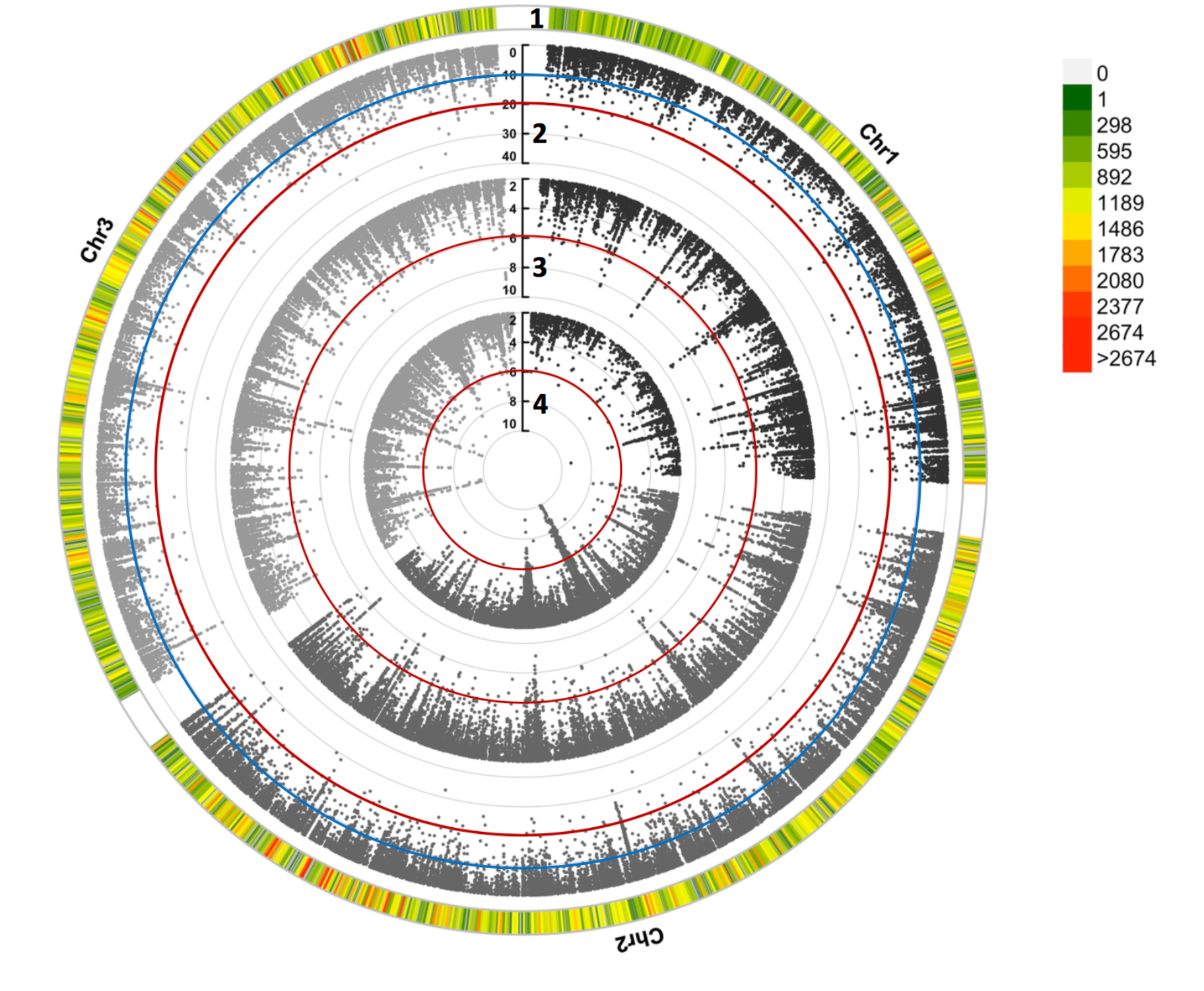
Circular Manhattan plot of genome-wide association analysis performed using three different methods. Ring 1 shows distribution of high-quality SNPs over three different *Ae. aegypti* chromosomes. It indicates the number of SNPs within 1 Mb windows and reflects the SNP density on each chromosome for genome-wide association with climate variables. Ring 2 shows SNPs association with MWMT (mean temperature of the warmest month) detected by BayPass. The significance level of association based on Bayes Factors (BF) is presented by blue (BF = 10) and red (BF = 20) circles. Rings 3 and 4 show SNPs detected by LFMM and PCAdapt respectively. The significance level (−logP for LFMM and PcAdapt) is represented by a red circle.

### Candidate gene functions and molecular pathways

We identified top candidate genes as those where an exceptional proportion of their total SNPs were outliers across all the three methods used, as explained above and in the methods section (Figure 4). In total, we found 112 top candidate genes and many of these genes were supported by multiple environmental variables (Supplementary Table 3). The vast majority of the genes detected as top candidates are annotated as being involved in a variety of biological processes, including AAEL001245 (EBgn0262737) which is known to encode a protein involved in thermo-sensory behavior and regulation of alternative splicing in *Drosophila* and AAEL019772 (FBgn0015245) and AAEL008641 (FBgn0001122) which encode heat shock proteins in *Drosophila* (56) (complete list of candidate genes with *Drosophila* homologs are described in Supplementary Table 3).

**Figure 4.**
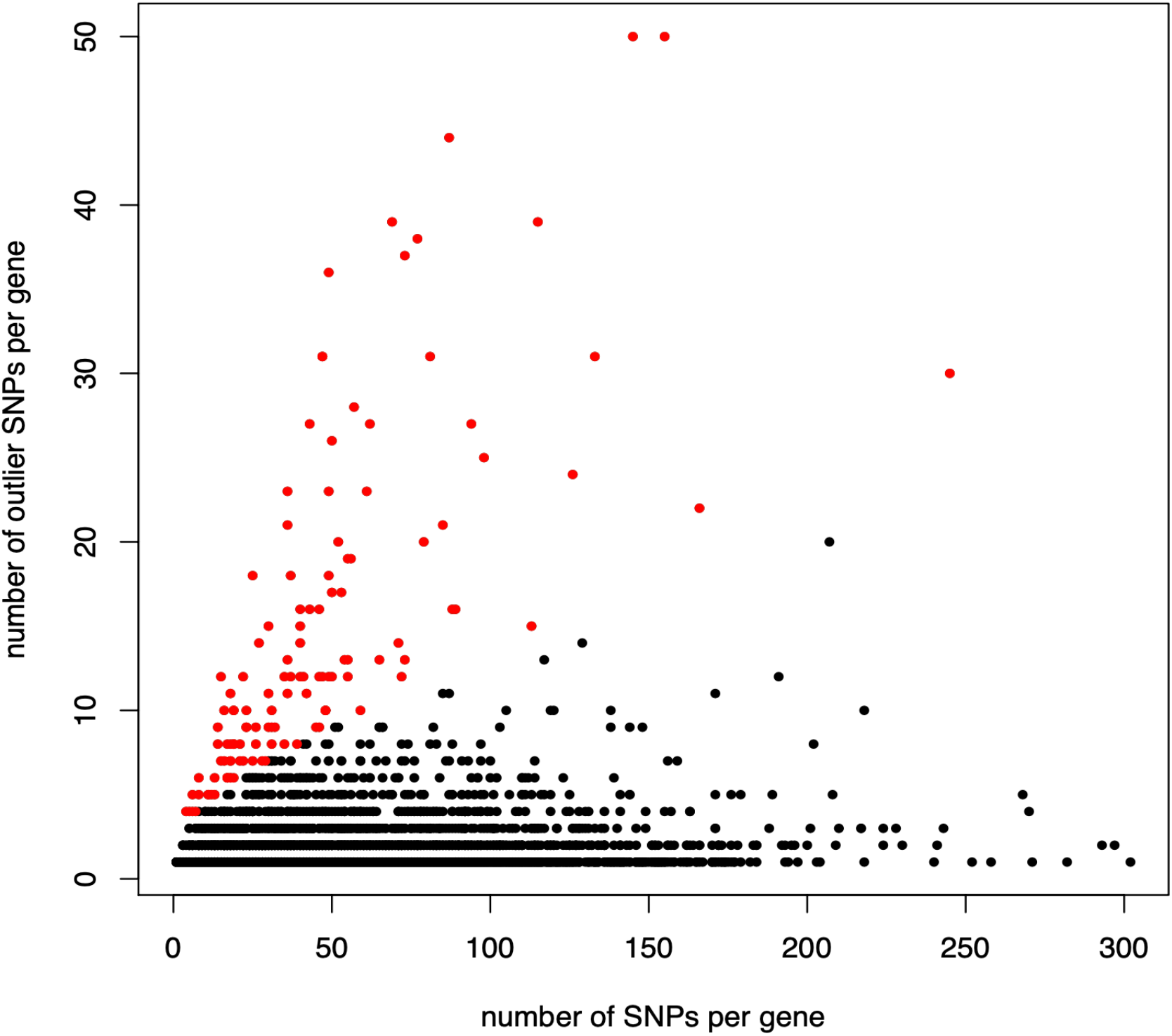
Top candidate genes for mean warmest month temperature (MWMT) identified as those with an extreme number of outlier SNPs relative to binomial expectation, shown in red. The same method was used to identify top candidate genes for each of the 25 topo-climate variable tests.

To understand the biological function of the top candidate genes, we performed GO enrichment analysis. From the 112 genes identified as top candidates, we identified 10 significantly overrepresented biological processes including metabolism, cell growth, response to stress, DNA repair, membrane assembly, transport through the endomembrane system, and mRNA transcription which all play important roles in adaption (Table 1).

**Table 1.** Top-ranked biological processes that were significantly overrepresented in the top candidate genes in *Ae. aegypti*.

| GO.ID | Term | Annotated | Significant | Expected | elimFisher | p.adj |
| --- | --- | --- | --- | --- | --- | --- |
| GO:0006468 | protein phosphorylation | 241 | 73 | 31.87 | 1.10E-12 | 0 |
| GO:0006355 | regulation of transcription | 403 | 100 | 53.29 | 4.70E-11 | 0 |
| GO:0007186 | G protein-coupled receptor signaling pathway | 151 | 46 | 19.97 | 1.70E-08 | 0 |
| GO:0035556 | Intracellular signal transduction | 184 | 55 | 24.33 | 5.70E-05 | 0 |
| GO:0035023 | Regulation of rho protein signal | 28 | 12 | 3.1 | 1.00E-04 | 0 |
| GO:0006813 | Potassium ion transport | 39 | 18 | 5.16 | 2.20E-04 | 0 |
| GO:0007165 | Signal transduction | 600 | 154 | 79.34 | 0.00028 | 0 |
| GO:0007169 | Transmembrane receptor protein tyrosine | 9 | 6 | 1.19 | 0.00031 | 0 |
| GO:0071805 | Potassium ion transmembrane transport | 13 | 7 | 1.72 | 0.00057 | 0 |
| GO:0007264 | Small GPTase mediated signal transduction | 17 | 25 | 9.52 | 0.00312 | 0 |

In order to gain further insight into the evolutionary history of adaptation, we performed nucleotide and differentiation-based tests to examine the presence of recent positive selection for the three genes with known activity in thermal adaptation (AAEL001245, AAEL019772 and AAEL008641). The nucleotide diversities (π) at the selected genes were significantly below the genome-wide averages in all three groups (Figure 5), which was consistent with the expectation of a strong selective event and rapid adaptive evolution (57). Additionally, the level of genetic differentiation (Fst) among populations was higher at the selected genes compared with genetic background, especially between *Ae. aegypti* 2 and *Ae. aegypti* 3 groups (Figure 5) implying that spatially varying selection has likely driven differentiation in these genes between the groups.

**Figure 5.**
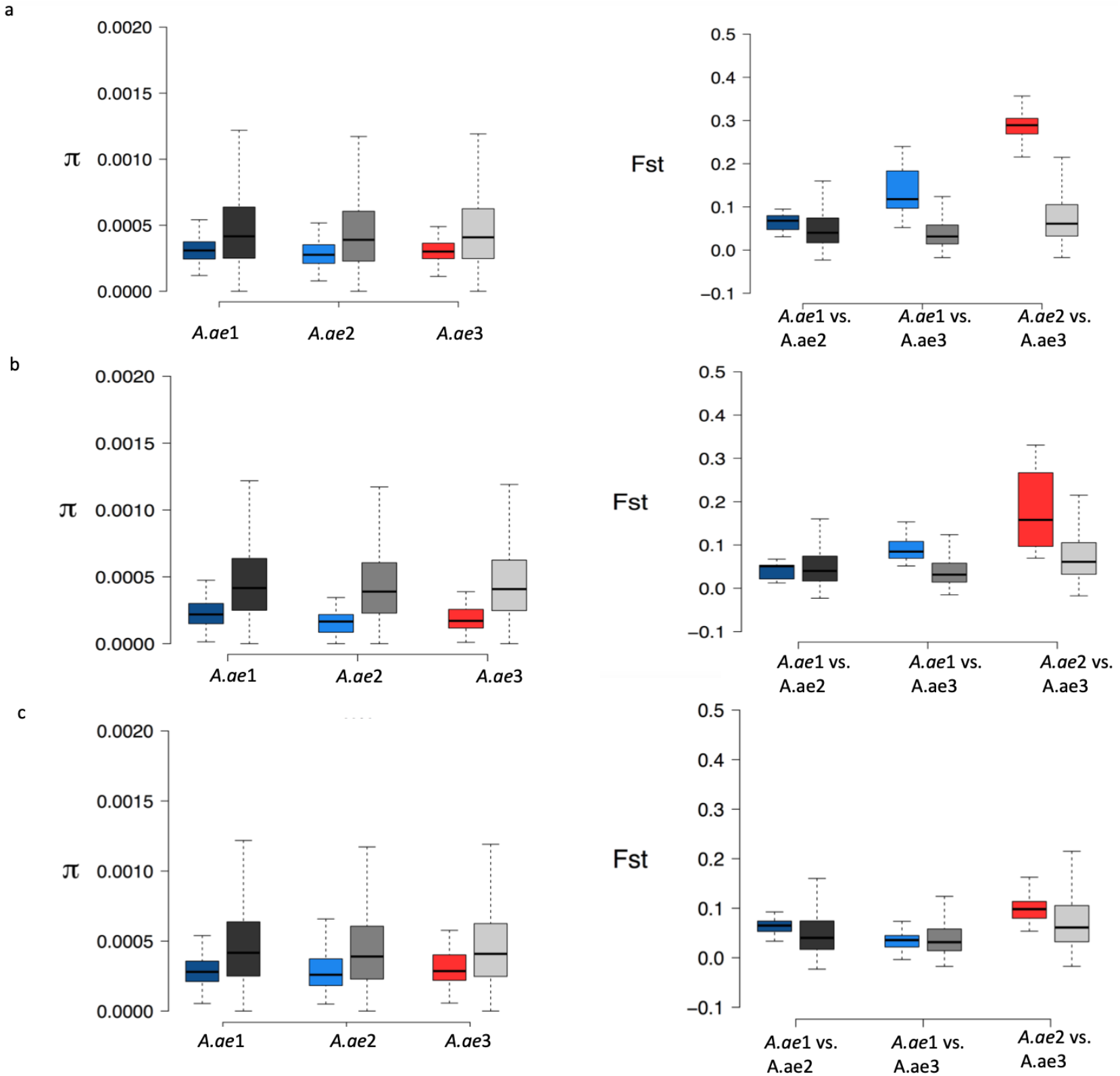
Nucleotide diversity (π) and genetic differentiation (Fst) for three genes, **a.** AAEL001245, **b**. AAEL019772 and c. AAEL008641, with significant signature of adaptation and well-known roles in thermal stress adaptation in *Drosophila melanogaster*.

## Discussion

Invasive species cause considerable ecological and economic harms worldwide (58, 59). Despite the broad impacts they have on diversity and agriculture, the genetic basis of adaptations and near-term evolution of invading populations are poorly understood. *Ae. aegypti* is the major vector of multiple diseases, such as dengue, Zika, and chikungunya and its geographical range is continuously growing; presumably due to anthropogenic conveyance, ongoing climate change, and increasing global transportation. The goal of the present study was to describe the fine-scale genomic architecture of this invasive mosquito within habitats characterized by different abiotic environmental conditions and to probe the underlying genetic basis for rapid adaptation of this species to new environments.

Our investigation of putative signals of selection and local adaptation of *Ae. aegypti* (in total 103 mosquitoes from 12 populations) that were recently introduced and established in various locations in central and southern California (Figure 1) found signals of selection, although not many, distributed along the genome. In a stepwise approach that included applying landscape genomics, identification of top candidate genes and GO enrichment analysis, we identified a set of candidate genes with various biological functions associated with adaptation to local abiotic environmental conditions in central and southern California.

Consistent with a previous study by Lee, Schmidt (18), we found structured populations of *A.aegypti* despite the potential for high levels of dispersal. We identified three major genetic clusters among the 12 populations collected across central and southern California, which seems to support the hypothesis that populations of *A.aegypti* distributed in California originated from multiple independent introductions from genetically distinct source populations; although the exact origin of the introductions remains uncertain and open for the future investigations.

To find genomic regions that have been targets of natural selection, we identified SNPs that are putatively selected for and strongly associated with topo-climatic variables using various landscape genomics methods. The methods we employed substantially controlled for neutral population genetic structure such as genetic drift or gene flow. We chose outlier SNPs as those which were consistently identified by all applied methods, allowing us to eliminate stochastic variation that could affect the results. We assumed that selected loci along the genome are likely to be under selection, either directly or through hitchhiking (60); although we acknowledge that other processes, including regions with reduced recombination, inversions, and chromosomal rearrangements, may also be responsible for the results we obtained (61). Therefore, further studies of linkage disequilibrium and the regions with reduced recombination and genome structure could illuminate the possible role of these factors in shaping adaptation as has been shown in other mosquitoes such as *Anopheles* (62).

Natural selection plays a key role in shaping the available genetic variation of populations and thereby produces adaptation (63). By applying EAA methodology we scanned the genome to uncover genomic selection footprints. We detected loci which were associated with both temperature and precipitation related variables (Supplementary Table 3), which implies the significance of both of these elements in shaping selection pressure and forming local adaptation in *Ae. aegypti*. Our results are in accordance with previous reports identifying these abiotic variables as major predictors in *Aedes* distribution patterns (14, 25). Temperature has been known to govern reproduction, maturation, and mortality rates and to be important for egg laying, development and survival of *Ae. aegypti* in larval habitats (64). These variables are also likely to elevate selection for thermal tolerance at the adult stage to resist diurnal and inter-seasonal variation (14, 64, 65). Precipitation affects the distribution of *Ae. aegypti* since rainfall generates breeding grounds for larvae and pupae. Unlike other mosquito species, *Ae. aegypti* eggs are laid above the water surface and hatch only when the water level rises and wets them (66).

We have identified signals of local environmental adaptation across a small number of large-effect loci and across a relatively moderate number of small-effect loci. Our lack of ability to detect more putative regions under selection can be explained by analytical limitations in distinguishing weak multi-locus signatures from the genomic differentiation introduced by genetic drift and demography (67, 68). In the presence of gene flow, selection is expected to act on a few number of loci with large effects since large effect loci are better able to resist the homogenizing effects of gene flow among populations (69). These limited regions are expected to have a strong impact on the fitness in one environment over the other because the allele with the highest fitness is expected to spread to all populations if this condition is not met. This can be tested in a common garden with a reciprocal transplant experiment in the future.

By applying top candidate gene method, we discovered 112 genes that contain SNPs highly associated with at least one topo-climatic variable (Supplementary Table 3). To better understand the role of each selected top candidate gene, we found their homolog genes in Drosophila. Several genes of heat shock protein (HSP) families are known to be selected in mosquitoes, which aid in overcoming stress induced by elevated temperature (70). Nucleotide diversity at these genes was below the genome-wide averages and the level of genetic differentiation was high among populations which further confirms that these genes are likely targets of the positive selection. In general, these results present some promising avenues for future works; especially if the markers detected here are linked to the actual targets of selection. The congruence between the observed genome scans and the genes assigned biological functions makes them ideal targets for further experimental validation. From an evolutionary perspective, coding regions are key genomic spots to look for the signatures of selection, as these directly influence functional elements in contrast to the non-coding genome regions. However, it is important to note that selection can also act on noncoding regions since they may be located, for example, in promoters, enhancers, or small RNAs where they affect gene expression. In this context, SNPs residing in non-coding regions of the genome may be of interest for future studies.

## Conclusions

The study of invasive adaptation and genome evolution is an emerging field that is developing rapidly and offers countless opportunities to investigate adaptive processes. Understanding the genomic basis of adaptive evolution in invasive species is important for predicting future invasion scenarios, identifying candidate genes involved in invasions, and, more generally, for understanding how populations can evolve rapidly in response to novel and changing environments.

Here we used a landscape genomics approaches to identify genomic regions and candidate genes potentially involved in adaptation. The identified genes showed footprints of selection and were correlated with environmental factors that differed between sites, as expected under a scenario of environment-mediated selection in natural populations of *Ae. aegypti* in California. Our findings help to elucidate the role of rapid evolution in the establishment and spread of invasive species. We detected evidence indicating local adaptation to various environmental conditions in populations of *Ae. aegypti* just a few years after its introduction into California, adaptations which may translate into a fitness advantage for specific populations.

## Funding

We acknowledge funding support from the Pacific Southwest Regional Center of Excellence for Vector-Borne Diseases funded by the U.S. Centers for Disease Control and Prevention (Cooperative Agreement 1U01CK000516).

## Acknowledgements

We thank personnel from Consolidated Mosquito Abatement District, Delta, Greater Los Angeles County, San Bernadino County, and San Mateo County Vector Control Districts and Coachella, Fresno, Madera County and Orange County Mosquito and Vector Control Districts, Community Health Division of the Department of Environmental Health, San Diego County Environmental Health, and Dr. Christopher Barker (UC Davis) for providing specimens used in this study. We thank Youki Yamasaki, Allison Chang, Parker Houston, Allison Weakley, Kendra Person, and Hans Gripkey for assisting DNA extraction and library preparations for this study. Thanks to Melika Hajkazemian, Melina Campos and Christine Coleman for their comments to the manuscript.

## Availability of data and materials

All scripts used for the analysis described are available on GitHub (https://github.com/shaghayeghsoudi/genomics_of_adaptation_A.aegypti_scripts) The datasets supporting the conclusions of this article are included within the article and its additional files.

## Author Contribution

ShS and GCL conceived the idea, designed the study, interpreted the data and wrote the manuscript. ShS performed the analysis and made figures. YL and AJC helped with gathering mosquitos. TCC performed alignment and variant calling. MC performed DNA extraction and library preparation. All authors read and approved the final version of manuscript.

## Competing interests

The authors declare that they have no competing interest.

## Notes

https://github.com/shaghayeghsoudi/genomics_of_adaptation_A.aegypti_scripts

### Competing Interest Statement

The authors have declared no competing interest.

